# Cyclic stretch inhibits cell invasion in 3D scaffolds

**DOI:** 10.64898/2026.06.13.732094

**Authors:** Rozanne Mungai, Juanyong Li, Jamie Baines, Leslie Kahugu, Kristen Billiar

**Affiliations:** Department of Biomedical Engineering, Worcester Polytechnic Institute, Worcester, MA 01605

**Author notes:** Correspondence to: Kristen L. Billiar, Biomedical Engineering Department Worcester Polytechnic Institute, 100 Institute Road, Worcester, MA 01609.

**Keywords:** mechanobiology, dynamic stretch, spheroid invasion assay, tissue engineered heart valves

## Abstract

**Background:** The development of clinically viable tissue-engineered heart valves (TEHVs) remains limited by inconsistent host cell infiltration. The dynamic hemodynamic environment may play a central role in driving or inhibiting cell invasion, yet the effects of cyclic stretch on cell migration and proliferation remain largely unexplored in 3D tissues and scaffolds. Given evidence that uniaxial constraint promotes directional invasion in 3D matrices, we hypothesized that uniaxial cyclic stretch would enhance cell invasion, particularly along the stretch direction.

**Methods:** We embedded multicellular spheroids into collagen hydrogels and subjected them to uniaxial cyclic stretch (3-10%, 1 Hz) for two days and quantified invasion into the surrounding extracellular matrix using a custom image-processing program. Smooth muscle cells, valvular interstitial cells, and dermal fibroblasts were examined to represent cell populations relevant to TEHVs and for comparison across cell types with different contractility. To determine the mechanisms underlying changes in invasion with stretch, effects of cell tension were evaluated using gel compaction assays and inhibition of myosin IIA, and proliferation was assessed by Ki67 immunostaining.

**Results:** Contrary to our hypothesis, cyclic stretch profoundly inhibited cell invasion into the matrix across all cell types and magnitudes of stretch. Invasion decreased by >50% in smooth muscle cells and fibroblasts and by up to 99% in valvular interstitial cells. Invasion suppression was inversely correlated with cell contractility, implicating a role for cell-generated tension. Inhibition of myosin IIA partially rescued invasion with stretch, though not to static levels. Stretched spheroids also exhibited reduced cell proliferation relative to static controls.

**Conclusions:** These findings implicate actomyosin-mediated mechanotransduction in stretch-induced suppression of cell invasion and suggest that the dynamic valve environment may limit host-cell repopulation of TEHVs. More broadly, this work provides insight into how cyclic stretch regulates 3D cell invasion in mechanically active tissues with implications for wound healing and cancer metastasis.

**Graphical Abstract:** 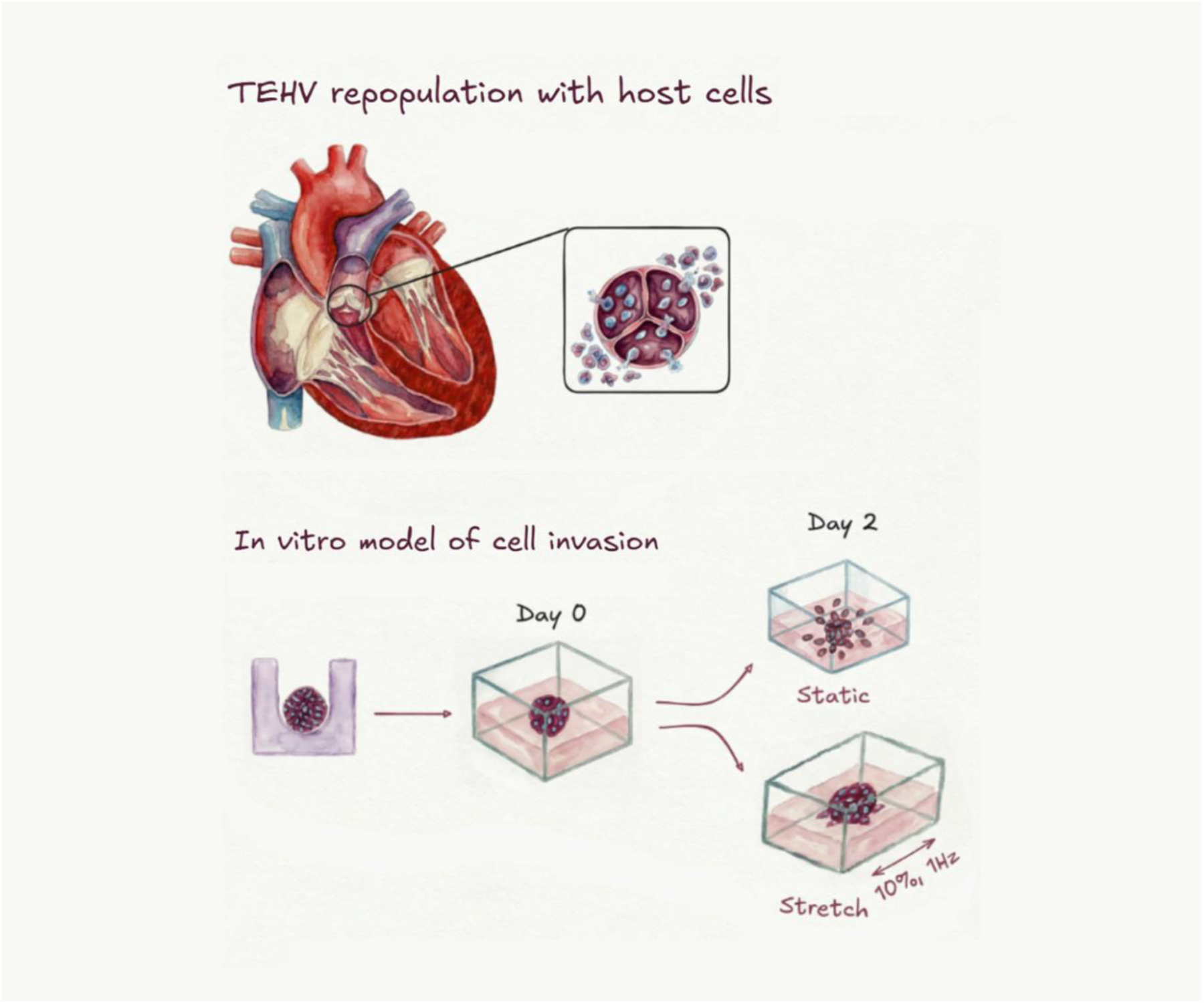

## INTRODUCTION

Tissue-engineered heart valves (TEHVs) represent a promising alternative to mechanical and bioprosthetic valves as they possess the potential to self-repair, adapt to patient-specific hemodynamic demands, and grow - features particularly critical for pediatric patients. Preclinical studies in sheep and primate models have demonstrated the feasibility of TEHVs composed of biologically engineered matrix in which engineered tissue is grown *in vitro*, decellularized, and implanted to allow repopulation and remodeling by host cells ^1–4^. However, cell repopulation remains largely confined to valve surfaces, with limited infiltration into the interior extracellular matrix (ECM), resulting in constructs with inconsistent growth and remodeling in the host ^2,5^. A deeper understanding of the mechanisms governing cell repopulation of the matrix is therefore critical for advancing TEHV design.

Cells attach and move within the ECM by interacting with the interconnected network of fibrous proteins. Cell–ECM adhesions act as mechanosensors, enabling cells to sense microarchitectural and mechanical cues in their environment ^6^ and migrate in response to mechanical stimuli, analogous to their chemotactic responses ^7^. It is also becoming increasingly clear that static mechanical cues such as stiffness ^8–11^ and contact guidance ^12,13^ play a large role in cell invasion ^7,14^. When presented with a steep stiffness gradient, fibroblasts accumulate preferentially towards stiffer regions on both 2D surfaces ^15^ and within 3D scaffolds ^16^. Similarly, invasive cancer cells preferentially migrate towards stiffer regions on 2D substrates where they exhibit increased cell traction forces indicative of elevated cell contractility ^17^. When cultured on aligned 2D topographies, cells undergo contact guidance that promotes directional migration ^7^. Similarly, in 3D collagen scaffolds, contact guidance governs directional invasion for a variety of cells such as fibroblasts ^13^ and cancer cells ^18,19^. In our work, we have shown that multicellular spheroids positioned near a stiff boundary undergo directional invasion towards the boundary, mediated by contact guidance generated through constraint-induced matrix remodeling and fiber alignment ^20^.

While the influence of static physical cues on cell invasion is well established, the effects of dynamic stretch remain largely unexplored, with most studies confined to 2D systems. In 2D contexts, cell migration responses vary with strain magnitude and frequency. Gefen and colleagues reported that fibroblast 2D migration increases with mechanical strain, with lower strain magnitudes (3% vs. 6%) ^21^ and slower stretch frequencies (0.1 Hz vs. 1 Hz) ^22^ producing the strongest responses. On the other hand, high magnitude stretch (10-15%) has been shown to reduce 2D cell migration of epithelial cells ^23,24^. Cyclic 10% stretch at 1Hz reduces cell migration of aortic smooth muscle cells in a scratch assay ^25^ and reduces invasion of bone marrow stromal cells that were pre-stretched prior to seeding on Matrigel-coated Transwells ^26^. Waters and colleagues have shown that 10% and 15% stretch at 0.16 Hz ^27^ and 20% stretch at 0.5 Hz ^28,29^ cyclic stretch decreases 2D migration of epithelial cells relative to static conditions. Collectively, these studies in 2D systems show inconsistent effects of cyclic stretch on cell migration, with responses varying by cell type, stretch magnitude and frequency. It remains unclear how these observations translate to 3D environments, where migration mechanisms differ and contact guidance within fibrillar matrices plays a dominant role.

Recent studies have revealed that cyclic stretch elicits changes in proliferation and contractility of cells, both of which are involved in cell invasion and migration. In 2D cultures, cyclic 10% stretch reduces cell proliferation in aortic smooth muscle cells ^25^. It also differentially regulates fibroblast cell contractility in a stiffness-dependent manner, increasing traction forces on soft substrates while reducing them on stiff substrates ^30^, and cell contractility has been inversely linked with cell motility, as partial inhibition of myosin II enhances cancer cell migration ^31^. Cyclic 10% stretch also induces fibroblast and mesenchymal stem cell elongation on 2D surfaces ^8,32^ and results in cell strain avoidance (i.e., cells align perpendicular to the direction of stretch) ^33,34–36^ for a variety of cell types such as endothelial cells, fibroblasts and cancer cells. In contrast, cells in 3D gels generally align along the direction of stretch, depending on fibrillation conditions and stretch timing ^33,37,38^. This distinction highlights that findings from 2D assays do not necessarily translate to 3D systems, underscoring the importance of 3D studies for understanding how mechanical cues govern cell behavior *in vivo*.

Although not dynamic, altering the mechanical boundary conditions of 3D model systems with static stretch, geometric constraint, and mechanical restraint has been shown to align ECM fibers resulting in enhanced cell migration via contact guidance ^18,39–41^. Mechanical boundary conditions can also increase the local apparent stiffness of the matrix and thus influence cell migration and invasion ^42,43^. Holmes and colleagues demonstrated that uniaxial restraint of a 3D hydrogel alters cell contractility, driving cells to generate anisotropic stress that promotes invasion from a cell-laden region into an acellular region along the axis of restraint ^42^. While mechanical boundary conditions in 3D systems promote directional invasion, whether these behaviors persist under dynamic stretch remains unresolved, motivating the need for further study.

We hypothesize that uniaxial cyclic strain enhances cell invasion in 3D scaffolds along the direction of stretch. To test this hypothesis, we embedded multicellular spheroids in collagen and fibrin hydrogels cast in stretchable silicone well plates and applied 3-10% cyclic strain at 1 Hz to the hydrogels for two days and quantified cell invasion using a custom image analysis program. Valvular interstitial cells and aortic smooth muscle cells were selected because they represent cell populations that may contribute to the repopulation of TEHV scaffolds *in vivo*. For comparison, dermal fibroblasts were included as a well-characterized fibroblast cell type, allowing broader interpretation of the findings and potential relevance to wound-healing processes. Multicellular spheroids were selected over other 3D invasion models such as cells coated over hydrogels for downward invasion ^44,45^ or nested cell-laden/acellular hydrogels ^42,46^ due to their ease of use and broad applicability across research contexts ^47–49^. The mechanisms of invasion were probed by inhibiting actomyosin contractility and quantifying proliferation.

## MATERIALS AND METHODS

There are no human or animal experiments in this study, and all cells were harvested from discarded tissues, therefore ethics approval is not required.

### Cell culture

Immortalized WKY 3M-22 male rat aortic smooth muscle cells (SMC) were obtained as a gift from Dr. Marsha Rolle ^50^. Porcine aortic valvular interstitial cells (VICs) were isolated from fresh male porcine aortic valves obtained from a local abattoir (Blood Farm, Groton, MA) within three hours of harvesting the hearts as per published protocols ^9^ and used in experiments between passages 2-8. Neonatal human dermal fibroblast cells (HDFs), harvested from de-identified donated male foreskins, were obtained as a gift from Dr. George Pins and used in experiments between passages 6-12. The same culture medium base formulation and conditions were used for all cell types: DMEM (Gibco) supplemented with 10% v/v fetal bovine serum and 1% v/v antibiotic-antimycotic (Gibco). For the SMCs, 1% MEM non-essential amino acids (Gibco) was added. The cell cultures were maintained at 37 °C in a humidified 10% CO_2_-containing incubator.

### Spheroid formation

Multicellular spheroids were generated for each cell type as per our previous work ^20^. Non-adherent agarose microwells were generated by pipetting a pre-warmed 2% w/v agarose solution (MilliporeSigma), made in DMEM, into negative molds with 500 μm diameter wells (Microtissues®), and the agarose was allowed to solidify. The cells were trypsinized and pre-stained with 5 μg/mL Hoechst 33342 (Invitrogen) for 10 minutes at 37 °C prior to spheroid formation. After dye dilution, the cells were resuspended at 1 × 10⁶ cells per 200 μL, seeded into the prepared agarose microwells, and allowed to settle into spheroids for one day before being harvested, resuspended into media, and embedded into collagen hydrogels.

### Preparation of hydrogels

For collagen gel experiments, cooled rat tail collagen type I (Advanced Biomatrix, RatCol®) was mixed with neutralization solution following manufacturer’s recommendations to make collagen hydrogels. For studies utilizing multicellular spheroids, a spheroid-media suspension was mixed into the collagen solution for ∼ 20 spheroids/mL in a final collagen concentration of 2 mg/mL. For proliferation studies utilizing single cells embedded in collagen, a cell suspension was mixed into the collagen solution at a final cell concentration of 0.5 x 10^6^ cells/mL.

For fibrin gel experiments, equal parts cooled 1.52 U/mL thrombin (Sigma Aldrich) dissolved in 60 μM calcium chloride (Sigma Aldrich) and spheroid media suspension (∼29 spheroids/mL) were combined and then mixed with an equal part of 8 mg/mL fibrinogen (Sigma Aldrich) ^51^ for final concentrations of 4.4 mg/mL fibrinogen, and 0.38 U/mL thrombin ^2^.

Next, either the collagen or fibrin gel solution was plated into the silicone well plate (4 x 4 array of 8 mm x 8 mm square wells) of a CellScale MechanoCulture FX™ uniaxial stretching device (Figure S1). The samples were placed into either the stretching device (dynamic condition) or a tissue culture plate (static condition) and allowed to gelate for either 30 minutes (collagen) or 1 hour (fibrin) at 37 °C. For a 2D control, some spheroids or cells (94,000 cells/mL) were also added directly into tissue culture dishes.

Fresh cell culture media was then added over the samples at the beginning of the two-day culture period. For experiments gauging how reducing cell tension affects cell invasion, para-amino-blebbistatin (Cayman Chemical) was added to the media at 0 μM, 2 μM or 20 μM. The samples were imaged (Day 0) prior to culturing for two days in a humidified 37 °C, 10% CO_2_-containing incubator at either a static or stretched condition (10% uniaxial stretch at 1 Hz frequency). For stretched samples, sterile water was added to the sacrificial wells to reduce media evaporation.

### Staining and Imaging

The extent of cell invasion from the spheroids into the surrounding collagen hydrogel was assessed by capturing z-stack images of the cell nuclei through the spheroid depth with a 10 µm step size as in our previous work ^20^. On Day 0 immediately after collagen gelation, the spheroids (pre-stained with Hoechst) were imaged live for comparison with images of the same spheroids on Day 2. At Day 2, the samples were fixed with 4% paraformaldehyde, permeabilized using 0.25% Triton-X 100 and stained again with Hoechst, at a 1:1000 dilution, to obtain a strong fluorescent nuclei signal. They were also stained for F-actin visualization using Alexa Fluor® 488 phalloidin (Life Technologies) at a 1:100 dilution. The images were captured using a Keyence BZ-X810 fluorescence microscope (6.1 µm DOF, High Resolution setting: 6 dB gain, Binning off) which allows for saving the Day 0 spheroid locations during imaging to ensure that the same spheroids were captured on Day 2.

The z-slices demonstrated consistent staining intensity through the depth of the spheroids ^20^. Spheroids that were out of the field-of-view or too close together (less than 400 µm apart or with invasion paths of different spheroids that could cross) were not imaged.

### Quantification of cell invasion into matrix

Maximum projections of the z-stacks and brightness and contrast pre-processing was performed using either the microscope software (Keyence BZ-X800 Analyzer ver 1.1.2.4) or Fiji (ver 1.54p, http://imagej.net) ^52^. Image quantification was carried out using a custom MATLAB program (ver. R2024a, MathWorks®, https://www.mathworks.com/) using the image processing package. The culture period of two days was chosen to allow sufficient time for cells to invade into the matrix while minimizing migration past the camera field of view. However, in the case of spheroids in the static condition and captured at 10X magnification, the invasion extent at times extended the field-of-view, so a circular mask was applied for those images to prevent directional bias due to the field-of-view and spheroids larger than ∼300 µm at Day 0 were omitted from analysis.

Image quantification of spheroid invasion was performed as in our previous work ^20^. In brief, the color images were converted to grayscale, contrast enhanced and binarized (Figure S2), using a user-defined global threshold, typically 0.16 from range [0,1]. The binarized Day 0 spheroid images were segmented to determine the spheroid centroid and the boundary of cell invasion. The centroids of the Day 0 and Day 2 image pairs were used to overlap the images to find the locations of all the pixels of Day 2 image that lay outside of the boundary of invasion (i.e., “the outer pixels”). The radial distances of each of these pixels from the boundary and the centroid were calculated as well as their angles of invasion.

The following metrics were then calculated as per our previous work ^20^. 1) The change in spheroid area *(ΔA)* from Day 0 to Day 2. 2) The mean distance per spheroid (*D̅*), calculated from the differences of the radial distance from each outer pixel to a corresponding point on the boundary. 3) The angles of invasion *(θ)*, calculated by the angle of each outer pixel measured clockwise from the x-axis. 4) The invasion area moment of inertia *(I)*, calculated by multiplying the squared radial or directional (x, y) distances (d) of each outer pixel *(i)* to the boundary with the area of each pixel and summing over all the outer pixels, Eq. (1). As explained in our previous work, *I* is an integrative measure of invasion which takes into account both the amount of nuclear staining (by the number of pixels) and the distances from the spheroid boundary. It allows for radial invasion quantification (*I_r_*) which is calculated using the radial (r) distances. It also allows for quantification of the directional moments of inertia which are the directions parallel and perpendicular to the direction of stretch (*I*_∥_ and *I*_⊥_) and are calculated using the directional (*x, y*) distances where *x* is the direction of stretch.

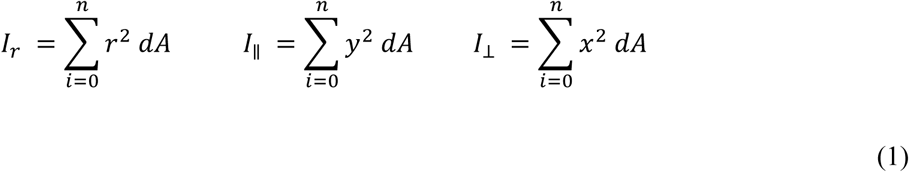

### Nuclear Alignment Analysis

To assess whether uniaxial stretch affects cell invasion direction and nuclear alignment, a subset of collagen-embedded SMC spheroids (stained with Hoechst and Phalloidin as above) were imaged using a Leica Stellaris 8 confocal microscope to acquire high-resolution images. Collagen fibers were also visualized using confocal reflectance at 488 nm. Z-stacks were captured through the spheroid depth with a 2.4 µm step size. Maximum projections of the z-stacks were generated using the microscope software and brightness and contrast pre-processing was performed using Fiji. The cell nuclei objects were segmented and identified using Cell Profiler (ver 4.2.6, http://www.cellprofiler.org) ^53^ using the IdentifyPrimaryObjects module with a global thresholding strategy and robust background thresholding method. Only individual (i.e., invaded) nuclei were analyzed by selecting objects with diameters between 3 and 20 pixels (scale ratio 1.14 µm/px); all objects outside this range were excluded from analysis. The eccentricity [0,1] and orientation angles [-90°, +90°] of the nuclei were quantified using the MeasureObjectSizeShape module and exported. Using MATLAB, histograms of the eccentricity and angles were plotted for each spheroid along with the corresponding probability density function (pdf). The nuclei angles (θ) and the probability distribution histograms of the nuclei angles, ℎ(θ), were used to calculate the orientation order parameter (S), Eq. (2), which is a single value [-1, +1] dictating the nuclei alignment for each spheroid ^54^). S = -1 and +1 indicate alignment parallel and perpendicular to stretch, respectively, while S = 0 indicates random or radial alignment.

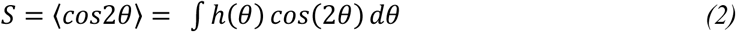

### Collagen compaction assay

To assess the differences in the tension that the three cell types generate in 3D culture, a standard collagen compaction assay was performed. Cooled rat tail collagen type I was mixed with neutralization solution as described above and then mixed with cells suspended in media for a final cell concentration of 1×10^6^ cells/mL in a final collagen concentration of 1 mg/mL. Next, the gel solution was plated into the wells of a non-adherent 24-well plate for a gel height of 2 mm and allowed to gel for 1-3 h at 37 °C. After gelation, photographs were captured of the gels in each well (0 h) using a Nikon stereoscope (SMZ-U) and they were detached from the sides of the wells using a metal spatula. Additional images were then captured to monitor cell compaction of the free-floating gels at 5, 10, and 30 minutes and 2, 4, 24 and 48 hours after detachment. The area of each gel at each timepoint was then measured using the polygon area selection tool in Fiji and was normalized to the initial area of the gel prior to detachment to calculate the percent area. The average area change was fitted to an exponential time decrease equation of a maxwell model using the MATLAB Curve Fitter tool with the nonlinear least squares method, without the robust method and with the Trust-Region algorithm. The time constant (τ), which describes the time required for the gel to reach 63% of its total compaction 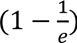, was obtained from the equation parameters. As a complimentary assay to our study of 3D cell contractility, traction force microscopy (TFM) was performed on individual cells in 2D culture to determine their contractility; the TFM methods and results (Figure S4) are included in the Supplementary Materials.

### Cell proliferation assay

To assess the effect of stretch on cell proliferation, we first attempted to stain and quantify Ki67-stained spheroids; however, the resulting signal was diffuse and difficult to threshold and segment using an automated approach (Figure S3). Therefore, we chose to embed SMCs as single cells in the collagen gels instead (300,000 cells/mL, 2.0 mg/mL collagen) and cultured them in stretch or static conditions for only one day to reduce cell-mediated gel compaction. After fixation, the 3D samples were stained with Hoechst (as above) and Ki67-AlexaFluor® 488 antibody (eBioscience) at a 1:250 dilution. Z-slices were captured using a Leica Stellaris 8 confocal microscope (range 160 µm, 10 µm step size) and processed into maximum projections. For 2D controls, 94,000 cells were seeded onto a 35-mm tissue culture dish and cultured for one day then fixed and stained with Ki67 antibody at a 1:500 dilution. Images were captured using the Keyence microscope. Brightness and contrast pre-processing for both 3D and 2D sample images were performed using Fiji. The nuclei density (i.e., Hoechst-stained cells) and fraction of nuclei expressing Ki67 (i.e., Ki67+ fraction) were quantified using Cell Profiler. In detail, the Ki67 objects were smoothed and closed to fill holes in the signal and the IdentifyPrimaryObjects module with a global thresholding strategy and robust background thresholding method was utilized to segment nuclei and Ki67 objects. The RelateObjects Module was then utilized to isolate the Ki67+ fraction of nuclei.

### Code availability

Our custom image analysis program, described in detail in our previous work ^20^, is shared on GitHub to facilitate use by other researchers and is available in MATLAB https://github.com/rmungai/SpheroidInvasionAnalysis as well as open-source Python https://github.com/rogerh2/SpheroidInvasionAnalysis versions with a downloadable GUI.

### Statistical analysis

Unless otherwise indicated, statistical analysis was performed using the sjstats library in R (ver. 4.4.3, https://r-project.org). An outlier analysis was performed on the dataset by detecting and removing values more than three scaled median absolute deviations (MAD) from the median. The normality of the data set was assessed via the Lilliefors normality test employing a significance level α=0.05. After determination of normality, significant differences among groups were analyzed. For two-group statistical analysis, unpaired Welch’s t-tests were used. For analysis of more groups, a one-way analysis of variance (ANOVA) was used, followed by Tukey-Kramer’s post hoc test. For determining the effect of stretch on invasion, the Cohen’s *d* effect size was calculated on the radial moment of inertia values for the static and stretch conditions as the difference between the group means divided by the pooled standard deviation. For non-normal data sets, two-group statistical analysis was performed using the Wilcoxon rank-sum test and the effect size was determined by the Wilcoxon effect size. The correlation between the compaction time constant (τ) and the effect size was plotted and calculated in Excel. Image quantification was performed with data pooled from 2-3 biological replicates, with at least three spheroids per experiment unless otherwise indicated. Violin plots were generated using the ggplot library in R to show the entire data distribution for all spheroids. Numbers of biological replicates (N) are provided in the figure captions as well as the numbers of spheroids or gels which are also indicated as dots in the plots.

## RESULTS

### Exploring the effect of stretch on cell invasion

We cultured multicellular spheroids embedded within collagen hydrogels for two days under static or uniaxial cyclic strain (3%, 10%, 1Hz) conditions to determine the effect of dynamic stretch on cell invasion. Contrary to our hypothesis, we observed a profound inhibition of invasion for stretched spheroids compared to the static condition for all three cell types, shown visually by a decreased nuclear staining beyond the initial spheroid boundary (Figure 1).

**Figure 1:**
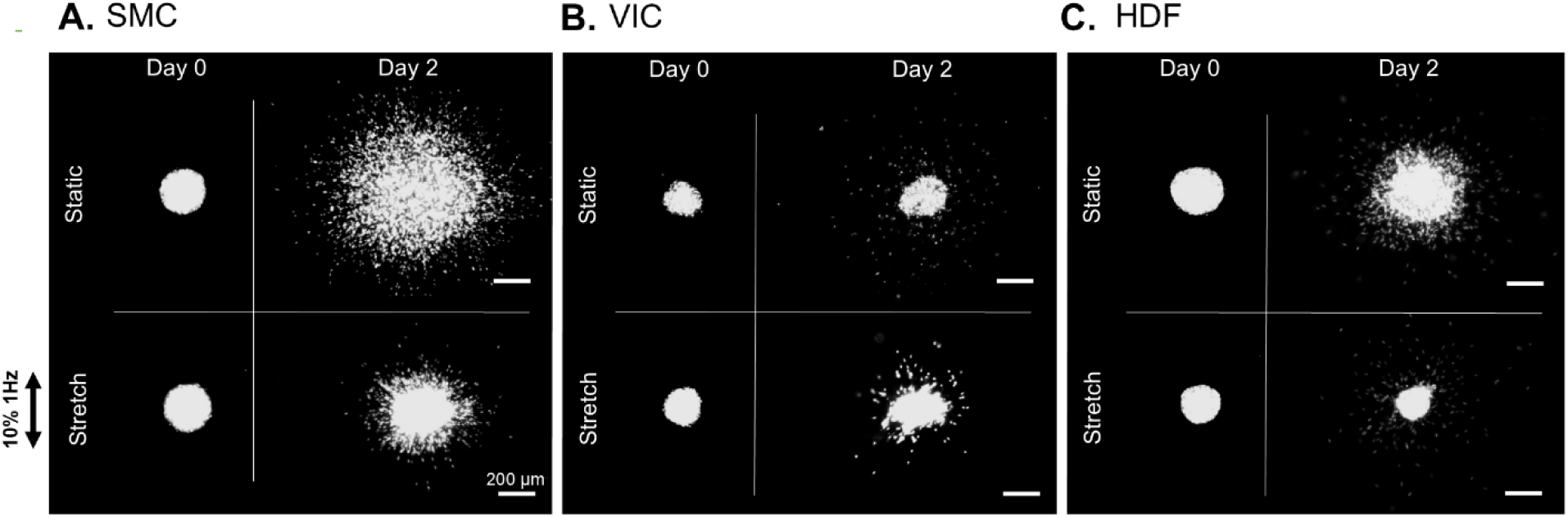
Cyclic stretch reduces cell invasion. Maximum projection images of cell nuclei of static versus stretched (10%, 1Hz) collagen-embedded multicellular spheroids on Day 0 compared to Day 2 for SMC (A), VIC (B), HDF (C). Grayscale images shown for clarity. Images captured at 10X magnification, scale bar: 200 μm. SMC spheroids subjected to 3% stretch invasion patterns to those at 10% and are therefore not shown.

The images then were binarized (Figure S2) and quantified to calculate the area change, mean distance, and radial area moment of inertia invasion (I_r_) metrics (Figure 2). Quantification confirmed that 10% uniaxial stretch at 1Hz significantly reduces cell invasion for all three cell types. Most strikingly, the radial moment metric (*I_r_*) reveals that stretch reduced invasion by over 50% for all cell types, and up to 90% for SMCs and VICs (Figure 2 A-C, Table S1B). However, the extent that stretch reduces cell invasion varies by cell type. The Cohen’s *d* effect sizes of the radial moments (stretch versus static) were calculated and found to be 2.40 for VIC, 2.01 for SMC, and 1.64 for HDF (Table S1A) indicating that, while stretch had a large effect on reducing invasion for all the cell types, SMCs and VICs exhibit substantially greater reductions of invasion than HDFs. We then tested whether the stretch-induced reduction of invasion is magnitude dependent by exposing SMC spheroids to a lower 3% uniaxial stretch at 1Hz. We observed that 3% stretch significantly reduced cell invasion by ∼90% reduction (Cohen’s *d*= 2.57; Figure 2D, Tables S1A, S1B). Notably, invasion at 3% stretch was not significantly different (p=0.804) from that at 10% stretch (Table S2).

**Figure 2:**
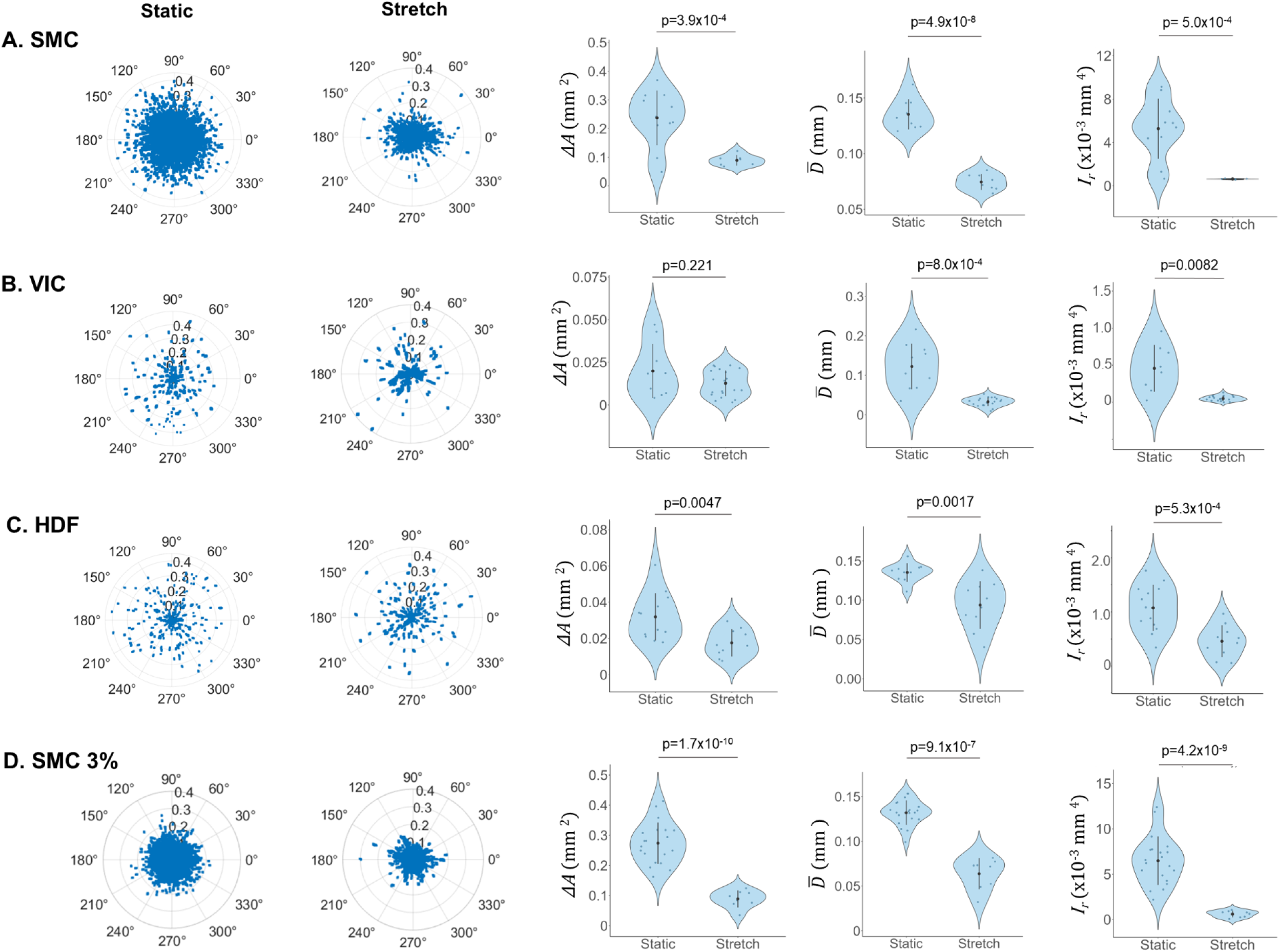
Cyclic stretch reduces cell invasion for SMC (A), VIC (B) and HDF (C) spheroids at 10% magnitude as well as for SMC spheroids at 3% magnitude (D). Polar plots depicting the invasion distances (mm) on Day 2 past the Day 0 boundary versus angle for all outer pixels of representative static and stretched spheroids (columns 1 & 2). Quantified metrics of cell invasion (columns 3 – 5): the area change (*ΔA*, mm^2^), mean distance (*D̅*, mm), and the radial moment of inertia (*I*_*r*_, mm^4^) confirm invasion reduction with stretch. Data are shown as mean ± SD. Number of spheroids (n), denoted as dots in the violin plots, and number of independent experiments (N) are as follows: static conditions, SMC (n = 9– 11, N = 2), VIC (n = 8–10, N = 3), HDF (n = 10–13, N = 2), and SMC 3% (n = 20, N = 1); stretch conditions, SMC (n = 5–8, N = 2), VIC (n = 19–21, N = 3), HDF (n = 10, N = 2), and SMC 3% (n = 8, N = 1). Statistical significance was assessed using an unpaired Welch’s t-test (p<0.05).

Unlike equibiaxial stretch, uniaxial stretch enables assessment of whether stretch direction influences invasion direction. We quantified the angles of invasion for each spheroid and calculated the directional area moment of inertia for the static and stretch conditions for each cell type (Figure 3). There was no significant difference observed for the directional moment values parallel or perpendicular to the direction of stretch (*I*_∥_ and *I*_⊥_ respectively) demonstrating that the direction of uniaxial stretch has no effect on invasion direction (Figure 3, Table S3). Similarly, there was no trend towards a particular direction for the angle histograms, rather cell invasion is radially orientated from the spheroid surface. In both static and stretched gels, we observed that individual invading cells displayed radially oriented F-actin fibers and nuclei, accompanied by local densification of the surrounding collagen matrix, consistent with active matrix engagement and remodeling (Figure 4A,B). Quantification of the eccentricity of the nuclei revealed distributions with a strong peak between 0.7 - 0.85 indicative of elongated nuclei characteristic of actively migrating cells (Figure 4C). In contrast, nuclear orientation exhibited more uniform distributions (Figure 4D), with orientation order parameter values near zero for both static and stretched conditions (static: 0.030, -0.106; stretch: -0.030, -0.090, -0.046). No significant differences in orientation were observed between conditions (Figure 4F, Table S4), indicating no evidence of directional bias in invasion. However, cell nuclei from stretched spheroids exhibited modestly reduced eccentricity compared with static controls (Figure 4E, Table S4), consistent with findings that reduced nuclear elongation correlates with reduced invasion ^55^.

**Figure 3:**
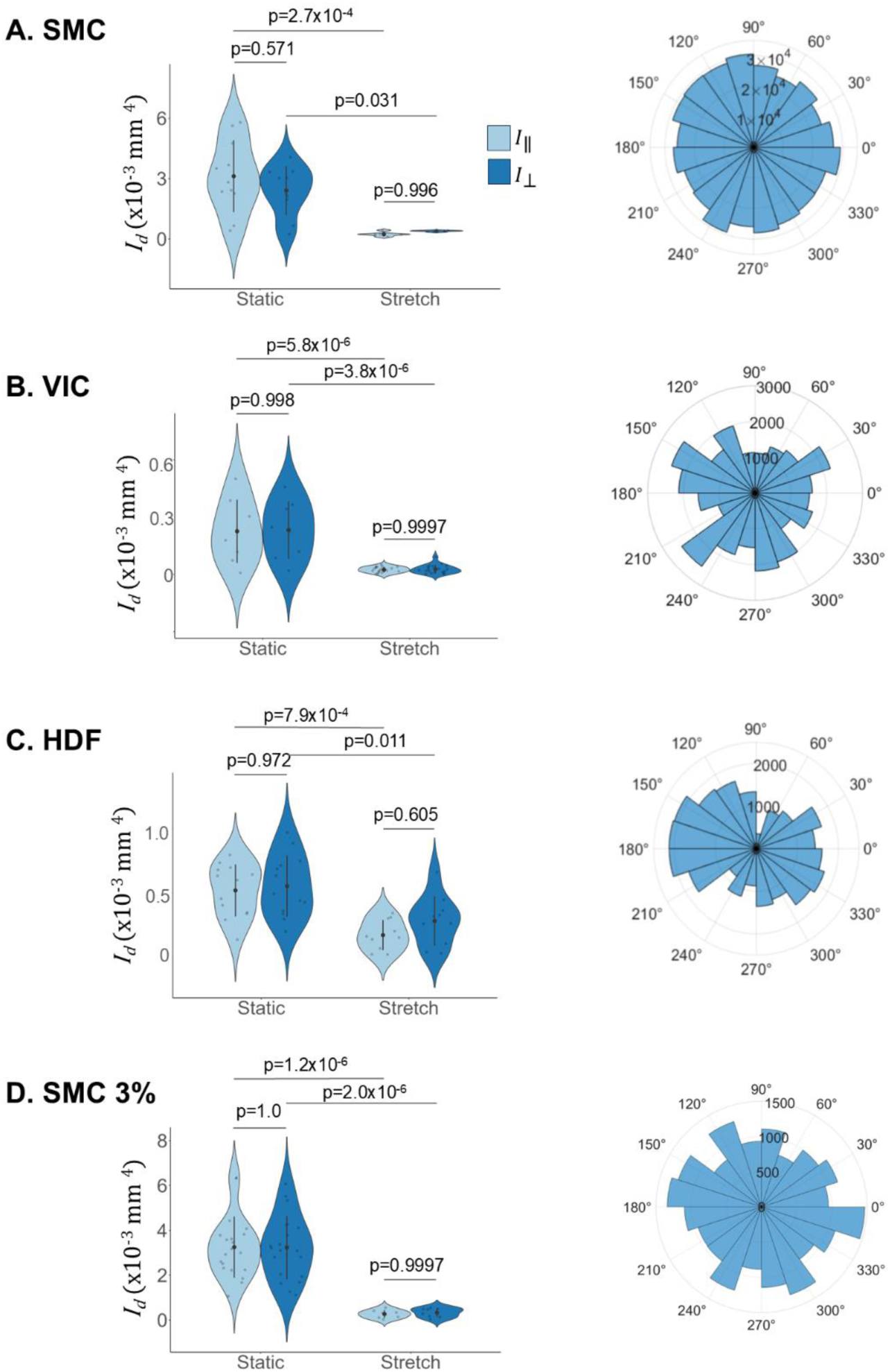
The direction of the uniaxial stretch does not influence cell invasion direction. Directional area moment of inertia *(I_d_*) for directions parallel *(I*_∥_) and perpendicular *(I*_⊥_) to stretch, and representative angle polar histograms for SMC (A), VIC (B) and HDF (C) spheroids stretched at 10% magnitude as well as SMC spheroids stretched at 3% magnitude (D). Data are presented as mean ± SD. Numbers of spheroids (n), denoted as dots in the violin plots and numbers of independent experiments (N) are as follows: static conditions, SMC (n = 11, N = 2), VIC (n = 8, N = 3), HDF (n = 13, N = 2), and SMC 3% (n = 20, N = 1); stretch conditions, SMC (n = 8, N = 1), VIC (n = 20, N = 3), HDF (n = 10, N = 2), and SMC 3% (n = 8, N = 1 Statistical significance was assessed using one way ANOVA with Tukey-Kramer’s post hoc test (p<0.05). P-values from all pairwise comparisons are included in Table S3.

**Figure 4:**
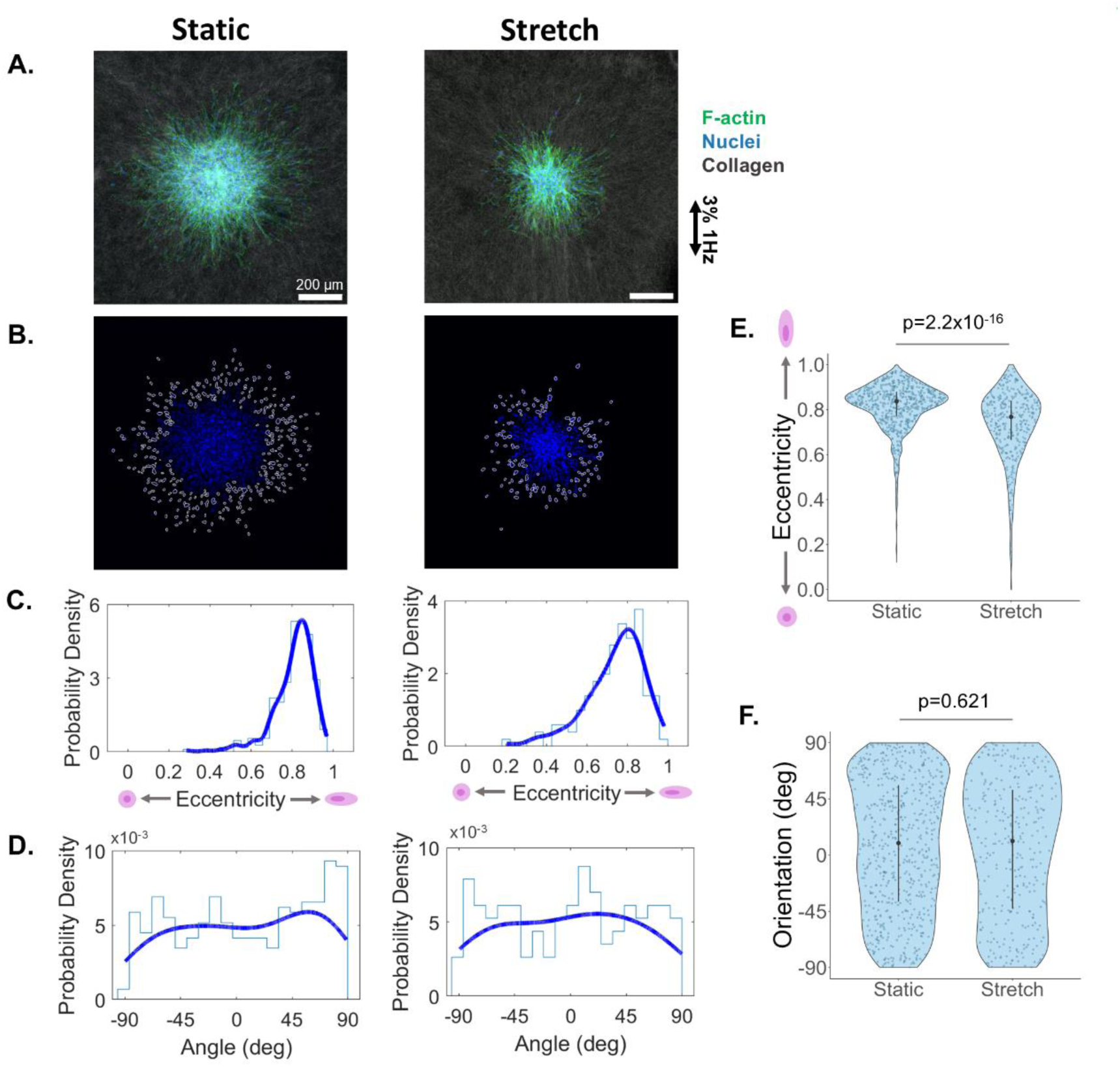
The direction of the uniaxial stretch does not influence nuclei orientation but reduces nuclei eccentricity. Representative maximum projection confocal images of static versus stretched SMC spheroids depicting cell F-actin fibers (green) and nuclei (blue) as well as the surrounding collagen fibers (gray) (A). Scale bar: 200 μm. Depiction of spheroid nuclei with individual invaded nuclei, as identified using Cell Profiler, outlined in white (B). The individual nuclei were quantified for the nuclear eccentricity [0,1] (C) and orientation angle [-90°, +90°] (D) with representative histograms (light blue) depicted along with the associated probability density functions (dark blue). Pooled eccentricity (E) and orientation values (F), data are shown as median and interquartile range due to non-normality. Numbers of cells, denoted as dots in the violin plots, are static = 631 cells from 2 spheroids and stretch = 339 cells from 3 spheroids. Statistical significance was assessed using the Wilcoxon rank-sum test (p<0.05).

### Exploring mechanisms of stretch-induced reduction of invasion

We next set out to determine the mechanism of the observed reduction of invasion for stretched spheroids. As the reduction was not stretch-direction dependent, we investigated active mechanisms. Because cell migration is strongly regulated by cell tension ^31^, we measured the contractility of individual cells using traction force microscopy and found that the most invasive cell type, SMCs (Figure 2A, D), are the least contractile (Figure S4). To extend this analysis to a 3D context, we used a collagen compaction assay as an indirect measure of contractility. Floating collagen matrices were generated for the three cell types and allowed to compact over two days. By comparing the rate of compaction between the cell types, we observed that HDF-seeded gels were the fastest to compact followed by SMCs and VICs, respectively (Figure 5A). This difference was especially evident at the 4-hour time point. Gel area was expressed as a percentage of the initial area at each time point and fit to an exponential time decrease curve to determine the compaction time constant (τ), defined as the time at which the gel reached 63% of its initial area (Figure 5B). To obtain a single representative τ value per cell type, curves were fit to the mean gel area values pooled across samples for each cell type. Consistent with these trends, HDF-seeded gels exhibited the lowest τ, followed by SMCs and VICs, respectively. The effect of stretch on spheroid invasion was quantified separately for each cell type as the Cohen’s *d* effect size of the radial moment metric, *I_r_* (Table S1A). These summary values, for which each cell type is represented by one single value (one τ and one effect size per cell type, n=3 cell types) were then correlated. Linear regression revealed a strong positive association between the I_r_ effect size and τ (R^2^ = 0.96) though the slope was not statistically significant with only three observations due to the pooled measurements (p=0.12) (Figure 5C). The HDF-seeded gels were the fastest to compact but were the least affected by stretch while the SMC- and VIC-seeded gels compacted the collagen slower and were more affected by stretch. These findings demonstrate an inverse relationship between cell contractility and stretch-induced invasion disruption.

**Figure 5:**
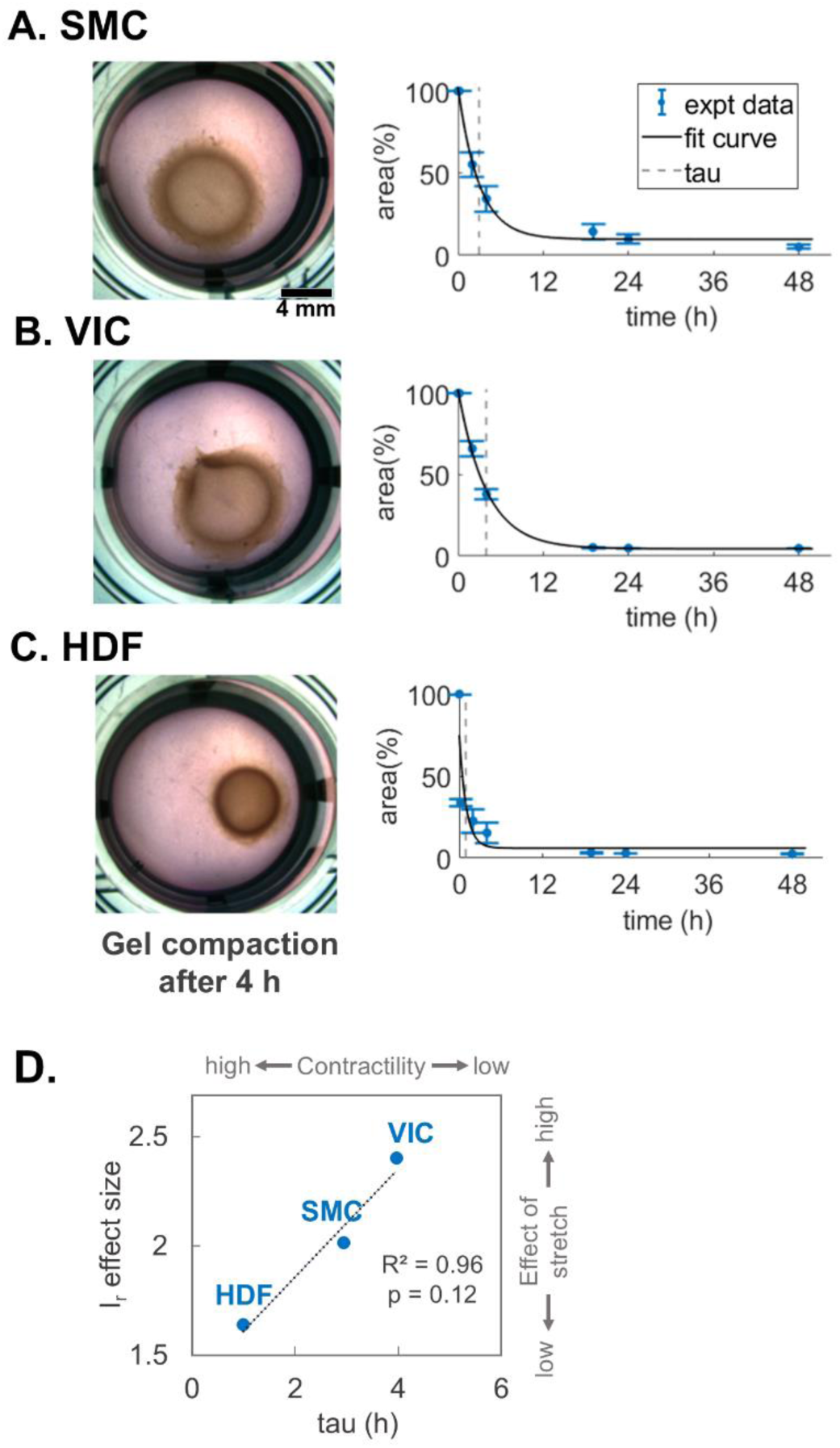
The effect of stretch is inversely related to the compaction time constant for the three cell types: SMC (A), VIC (B) and HDF (C). Representative images of collagen gel compaction after 4 h (left column). Scale bar: 4 mm. Quantified gel percent area relative to initial area (mean ± SD shown) fitted to an exponential time decrease curve (right column). The time constant (τ), representing the time at which the gel was 63% of its initial size, is indicated by a dashed line. Correlation plot of the effect of stretch on spheroid invasion and the time constant (D). A linear regression was fit to the data (y = 0.2475x + 1.3626; R^2^ = 0.9647), with statistical significance defined as p<0.05. Slope p-value: p = 0.12, 95% CI: [-0.35, 0.85], n=3. The effect of stretch is quantified as Cohen’s *d* effect size, based on the radial moment of inertia (*I_r_*) for stretched versus static spheroids. Numbers of free-floating gels are: VIC n=4 hydrogels, HDF, SMC n=5 hydrogels from one independent experiment. Numbers of spheroids and corresponding numbers of experiments is consistent with earlier figures.

Cells compact collagen gels by exerting cytoskeletal contractile forces that pull on collagen fibers. Since we observed an inverse trend between cell contractility (indicated by rate of gel compaction) and the effect of stretch on invasion, we then directly explored whether increased cell tension could be the mechanism underlying reduced invasion due to stretch. We treated SMC multicellular spheroids with a low dose of blebbistatin to reduce cell tension by inhibiting myosin IIA; this treatment has been shown to increase 2D cell migration ^56^. We found that 20 μM blebbistatin, which results in the characteristic shrunken cell morphology in 2D control cultures (Figure 6B), significantly decreases invasion in static spheroids (Figure S5), indicative of inhibition of cell contractility. Paradoxically, in stretched spheroids, 20 μM blebbistatin increases 3D cell invasion (Figure 6A, C, Figure S6). Yet even with this increase, the invasion (*I_r_*) is still 70% lower than for the control static group (Table S5) indicating that blebbistatin only partially rescues invasion in the stretched gels (Figure 6C).

**Figure 6:**
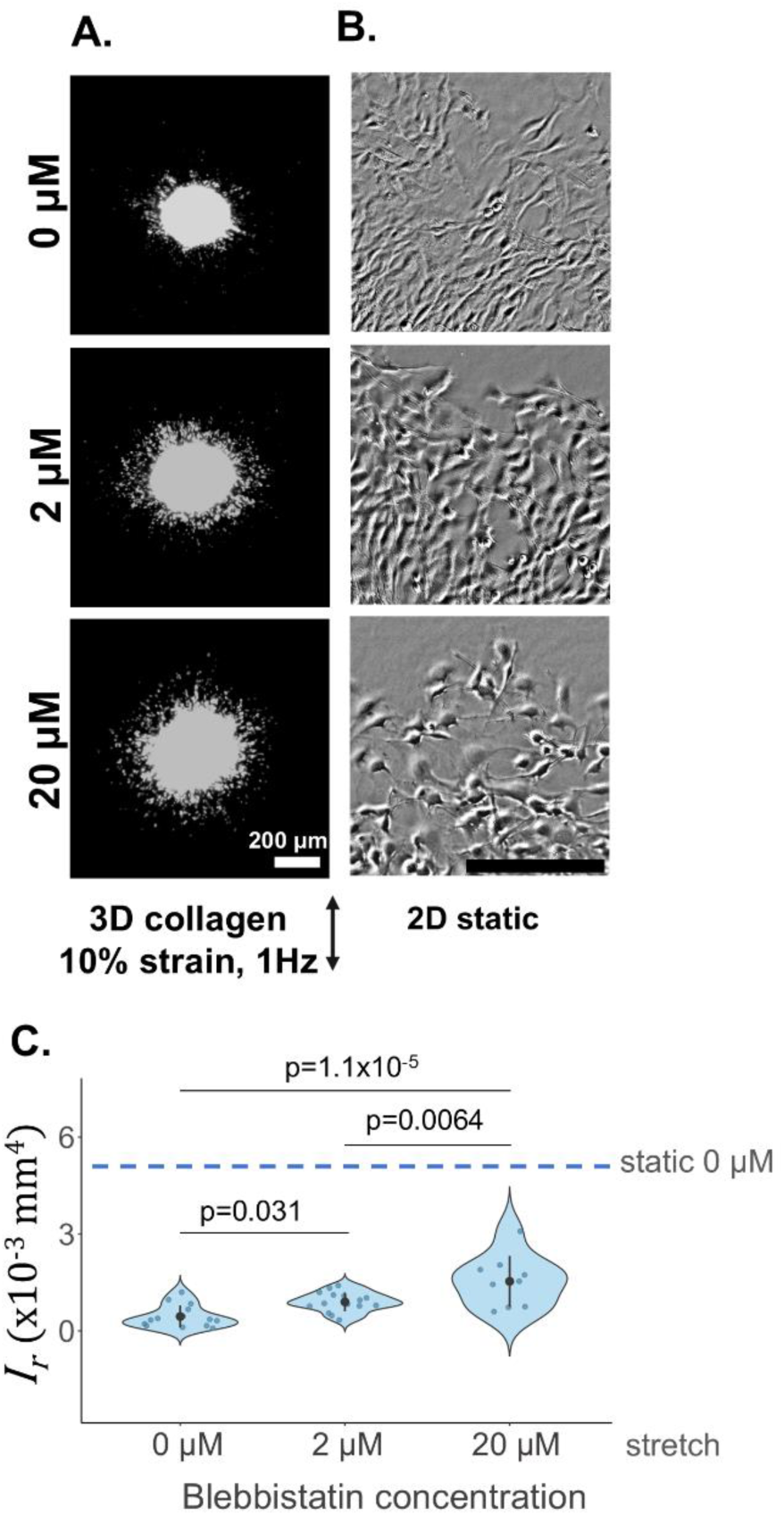
Blebbistatin partially rescues stretch-induced invasion reduction. Representative Hoechst-stained SMC spheroids (grayscale) showing 3D invasion under 10% uniaxial cyclic stretch (A) and phase-imaged SMC spheroids seeded on 2D tissue culture plastic illustrating cell morphology under treatment with 0 µM, 2 µM and 20 µM blebbistatin (B). Scale bar, 200 µm. Quantified area moment of inertia (*I*_*r*_, mm^4^) (C). Data are shown as mean ± SD. Dotted line denotes mean from statically cultured SMC spheroids untreated with blebbistatin (Figure 2A). Numbers of spheroids (n), denoted as dots in the violin plots, and numbers of independent experiments (N) are as follows: stretch 0 µM (n = 14, N = 3), stretch 2 µM (n = 16, N = 3), stretch 20 µM (n = 9, N = 1), and static (n = 10, N = 2). Statistical significance was assessed using one-way ANOVA with Tukey-Kramer’s post hoc test (p<0.05).

Cell proliferation has been shown to play a role in cell invasion of various cell types ^57–59^; thus, we explored whether stretch-induced changes in proliferation could be, in part, responsible for effects of stretch we observed. To determine if the invasion reduction under 10% stretch is due to decreased cell proliferation, SMC-embedded collagen gels were either stretched or kept in a static condition for one day before they were fixed and stained for the Ki67 proliferation marker. We found that 10% cyclic stretch reduced cell proliferation as evidenced by a reduced number of nuclei and reduced Ki67 signal (Figure 7). In 3D collagen gels, stretch reduced the number of cell nuclei by 20% and the fraction of Ki67+ cells by 40%, indicating that stretch reduces cell invasion partially by reducing proliferation (Figure S7 C). Compared to static 2D culture, SMCs grown in static 3D collagen were less proliferative and expressed 80% fewer cell nuclei and 35% fewer Ki67+ cells (Figure S8).

**Figure 7:**
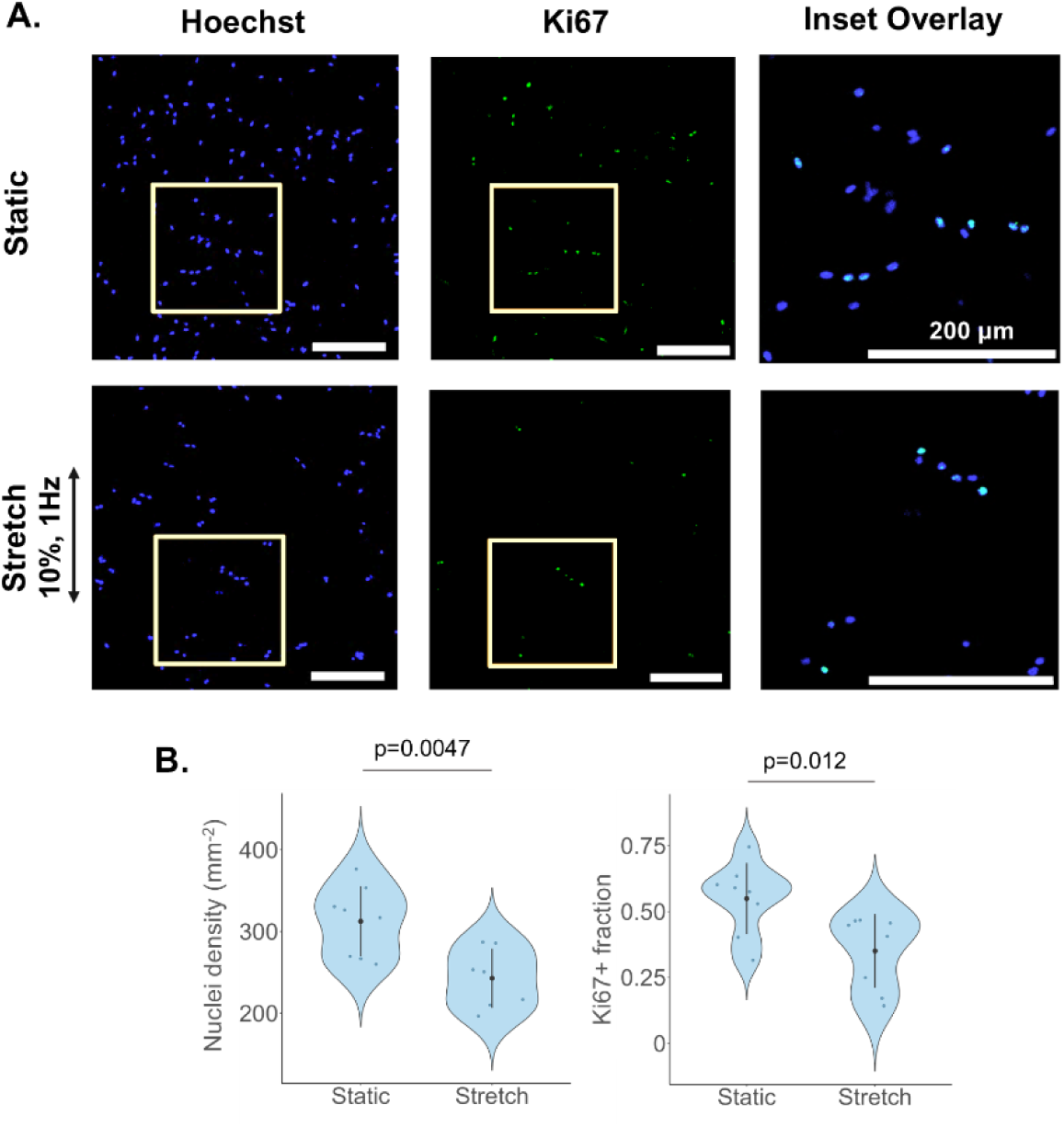
10% cyclic stretch reduces 3D cell proliferation. Representative images of SMCs embedded in static or stretched collagen hydrogels stained for the Ki67 proliferation marker (green) and Hoechst for cell nuclei (A). Inset denotes magnified region to visualize Ki67+ cell fraction expressing both Ki67 and Hoechst (overlay image). Scale bar 200 µm. Quantified nuclei density from Hoechst+ cells and fraction of Ki67+ cells for static and stretched samples (B). Data are shown as mean ± SD. Numbers of gels are: nuclei density static n=8 and stretch n=7, Ki67 static and stretch n=8. Data shown from one representative experiment, with a second independent experiment demonstrating similar trends provided in Figure S7. Statistical significance was assessed using an unpaired Welch’s t-test (p<0.05).

We next examined whether the stretch-induced reduction in invasion depends on ECM composition. SMC spheroids were embedded in fibrin hydrogels and cultured for two days under either static conditions or cyclic stretch (10%, 1 Hz). As observed in collagen gels, stretch reduced cell invasion in fibrin hydrogels across all invasion metrics (Figure 8A–E, Figure S9) and, in particular, decreased the invasion moment (*I*_*r*_) by 78%, corresponding to a large effect size (Cohen’s *d* = 2.04; Table S6). Comparing collagen and fibrin matrices, we observed that under stretched conditions, invasion was higher in collagen than in fibrin (46% higher *I_r_*; Figure 8F) with a large effect size (Cohen’s *d*=1.46; Table S6), although this difference did not reach statistical significance.

**Figure 8:**
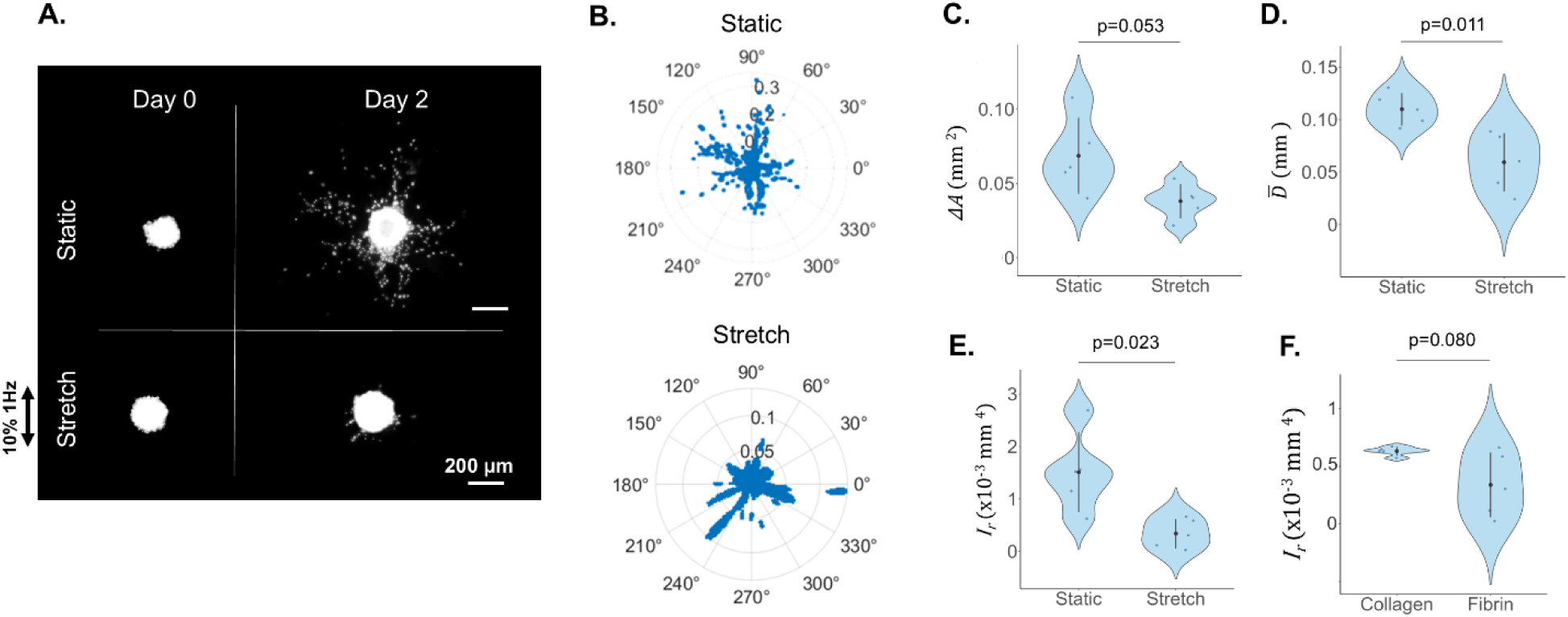
10% cyclic stretch reduces 3D cell invasion in fibrin gels. Representative images of Hoechst-stained SMC spheroids showing 3D invasion under 10% uniaxial cyclic stretch (A). Grayscale images shown for clarity. Scale 200 µm. Polar plots of invasion distances for representative static and stretched spheroids. Plot depicts distances (mm) versus angle for pixels extending past the Day 0 boundary (B). Note different axes for static and stretch cases. Quantified area change (ΔA, mm^2^), mean distance (*D̅*, mm), and radial moment of inertia (*I*_*r*_, mm^4^) for static versus stretched SMC spheroids in fibrin gels (C - E). Radial moment of inertia for stretched SMC spheroids embedded in collagen versus fibrin hydrogels (F). Data are shown as mean ± SD. Numbers of SMC spheroids in fibrin gels, denoted as dots in the violin plots, are n=5 for N=1 experiment for both stretch and static conditions. Statistical significance was assessed using an unpaired Welch’s t-test (p<0.05).

## DISCUSSION

In this work, we hypothesized that uniaxial cyclic stretch would enhance cell invasion into extracellular matrices along the direction of stretch based on findings that stiff boundaries increase cell invasion into soft scaffolds along the direction of constraint. However, we found that cyclic stretch sharply decreases cell invasion from spheroids embedded within both collagen and fibrin hydrogels, although the extent that stretch reduces invasion differs by cell type. Contrary to our hypothesis, the direction of uniaxial stretch does not influence the decrease in invasion, and inhibition of invasion occurs equally at low (3%) and high magnitudes (10%) of stretch. Investigations into the underlying causes suggest that cyclic stretch suppresses invasion through at least two contributing mechanisms: reduced cell proliferation, indicated by lower Ki67 levels, and elevated cell tension, indicated by the partial rescue of invasion with blebbistatin. Yet these factors account for only moderate inhibitory effects on invasion suggesting that additional unidentified mechanisms also contribute to the profound decrease in invasion induced by stretch.

### Stretch-induced suppression of invasion

Despite the mechanically dynamic nature of many tissues, the effect of stretch on cell migration or invasion has scarcely been studied. Using our multicellular spheroid-based *in vitro* model, we found that 10% uniaxial cyclic stretch reduces cell invasion into 3D matrices compared to static conditions (∼90% reduction; Figure 2). This finding is consistent with the limited observations that have been made in 2D dynamic scratch assay studies where 10-20% cyclic stretch leads to reduced migration into the wound gap ^25,27,60^. A similar effect has been reported in a 3D wound healing model, in which 2 – 2.5% cyclic stretch reduced dermal fibroblast infiltration from a cell-laden collagen gel into a nested acellular gel ^61^. Notably, our findings further suggest that stretch-induced inhibition of invasion is largely independent of stretch magnitude (3% versus 10%), in contrast to the broader 2D literature, where responses are highly context-dependent: low 3% stretch promotes cell migration in fibroblasts ^21,22^, moderate 5% stretch at a very low rate (0.16 Hz) has no effect in epithelial cells ^27^ and high 15-20% stretch reduces migration in epithelial _cells_ ^27,29,62,63^.

### The effect of stretch direction

In 3D hydrogels, anisotropic uniaxial mechanical boundary conditions such as static stretch ^64^, geometric constraint ^65^ and mechanical restraint ^42^ promote cell reorientation, leading to preferential cell alignment ^65^ and directional invasion along a defined mechanical axis through ECM fiber alignment and contact guidance ^39,42,64^. Based on these findings, we hypothesized that uniaxial stretch would similarly align collagen fibers and cells along the stretch axis, thereby enhancing invasion along the direction of stretch. Contrary to this expectation, we did not observe a directional bias in invasion for any of the three cell types (Figure 3). Alignment of collagen fibers along the direction of low magnitude (2%) uniaxial cyclic stretch has been reported, yet even with this alignment no preferential migration in this direction was observed, consistent with our finding ^61^. The lack of directional bias in our results may arise from the high cell density within multicellular spheroids. Cells are known to polarize away from regions of high density, leading to radial alignment at the spheroid surface prior to invasion ^66–69^. As a result, cells may preferentially migrate away from the dense core rather than respond to externally applied mechanical cues ^70^. Consistent with this interpretation, our confocal imaging showed that invading cells interacted with an unaligned collagen matrix while maintaining predominantly radial nuclear orientations, rather than aligning parallel or perpendicular to the stretch axis (Figure 4). Thus, intrinsic radial organization appears to dominate over applied stretch, resulting in radial rather than stretch-guided invasion, even far away from the spheroid. It remains possible that stretch-guided orientation would emerge in alternative 3D models with lower cell density (i.e., non-spheroid systems).

### The role of matrix composition

It has long been established that the matrix protein composition strongly influences both the rate and mode of cell migration. As early as 1993, Dvorak and colleagues observed that fibroblasts are six times less migratory in fibrin compared to collagen hydrogels ^71^. Similarly, in our gels of similar composition, we observed approximately half the extent of cell invasion in fibrin compared to collagen hydrogels for both static and stretch conditions (cf. Figures 2A, 8B-E), and invasion of stretched spheroids appeared more extensive in collagen than in fibrin, although this difference did not reach statistical significance (Figure 8F, Tables S1, S6). In addition, we have previously shown that fibroblasts spread 40% more over collagen than fibrin hydrogels and sense mechanical constraints 30% further through collagen than fibrin gels ^72^. Since cell spreading and adhesion are key components of mesenchymal cell migration ^73^, the greater spreading observed in collagen gels compared with fibrin may contribute to the more extensive invasion in collagen gels. Further, supplementing collagen with other matrix proteins has been shown to enhance invasion; for example, collagen I matrices supplemented with fibronectin, tenascin-C and collagen IV significantly increase 3D spheroid invasion compared with collagen I alone in static culture conditions ^74^. Similarly, hydrogels containing both collagen and fibronectin promote invasion during magnetic bead-induced twisting, whereas collagen-only gels show a slight, non-significant reduction of invasion ^75,76^. Although matrix composition strongly influences invasion, we observed no significant difference between fibrin and collagen matrices, suggesting that stretch-induced invasion reduction is largely independent of matrix composition and not primarily governed by cell-matrix adhesion. This finding instead points to a cell-intrinsic mechanism, such as cell tension.

### The role of cell tension

Cell tension has been implicated in regulating cellular invasiveness, with studies reporting both positive and negative relationships between contractility and invasion depending on context. Increased invasion has been associated with higher contractility in some 3D cancer models ^17,77^, but with lower contractility in 2D migration of stem cells ^56^ and liver pericytes ^78^, as well as in other 3D cancer models ^31,79^. Given these seemingly conflicting relationships, we investigated whether differences in contractility could explain the variation observed in both invasiveness and stretch-induced inhibition of invasion across the three cell types, using traction force microscopy and a collagen gel compaction assay. Although cell compaction of collagen gels is not a direct measure of cell contractile force, biochemical modulators of contractility (e.g., blebbistatin, LPA, TGFβ) influence compaction rates under controlled conditions ^80–83^. Here we show that faster gel compaction (lower τ) is associated with a reduced effect of stretch on invasion (Figure 5), indicating that highly contractile cells are less sensitive to stretch-induced inhibition. We also show that SMCs, the most invasive cell type, exhibit the lowest contractility by TFM. Although these single-cell measurements do not correlate with the effect of stretch on invasion across the cell types, likely due to differences between individual (2D) and collective (3D) behavior as well as contributions from factors such as proliferation, they nevertheless support an inverse relationship between invasiveness and contractility in a cell-type-dependent manner.

As cell contractility is primarily driven by actomyosin interactions, we then explored the role of myosin IIA on stretch-induced invasion reduction. Because myosin IIA expression increases with collagen stiffness ^84^ and mechanical stretch increases the local stiffness of hydrogel fibers ^85^, we tested whether cyclic stretch enhances myosin IIA–mediated tension by treating spheroids under static and stretched conditions with low-doses of blebbistatin. Blebbistatin has been shown to increase cell migration on 2D substrates ^56,78^, decrease cell migration in 3D collagen ^83^ and fibrin hydrogels ^86^ and reduce gel compaction in free-floating 3D collagen hydrogels in a dose-dependent manner ^56,80,83,84,87^. We found that increasing blebbistatin concentration enhances cell invasion from stretched spheroids (Figure 6); however, static spheroids show a non-monotonic response, with invasion increasing at 2 μM but decreasing at 20 μM blebbistatin (Figure S5). This difference shows that optimal motility requires tensional homeostasis ^88^, characterized by balanced myosin IIA activity. In static conditions, partial inhibition of myosin IIA can promote motility, while stronger inhibition reduces motility. In contrast, stretch upregulates myosin IIA and elevates cell tension, shifting contractility out of balance and limiting invasion, and as a result, higher blebbistatin concentrations are required to restore balance. Thus, with stretch, 20 μM blebbistatin is sufficient to partially rescue invasion whereas this high dose suppresses invasion in static conditions. Further, actomyosin-generated tension directly influences nuclear morphology, with increased cellular tension leading to nuclear elongation or deformation ^89,90^. Since migrating cells generally have elongated nuclei with higher eccentricity ^55,91,92^, our observation that stretch decreases nuclear eccentricity (Figure 4) further supports a role for actomyosin contractility in mediating the reduced invasion observed with stretch. Together with the collagen compaction results, these findings implicate actomyosin contractility as a contributor to inhibition of invasion by cyclic stretch.

### The role of cell proliferation

In addition to cell contractility, cell proliferation also contributes to cell invasion from multicellular spheroids ^57–59,93^. Proliferation has been shown to be modulated by dynamic stretch, either increasing ^94^ or decreasing ^25^ depending on stretch magnitude, frequency and duration ^95^. Here, we have shown that 10%, 1Hz cyclic stretch reduces cell proliferation in 3D collagen gels (Figure 7). Our findings are consistent with prior 2D scratch assay studies in which SMCs exposed to 10%, 1Hz uniaxial stretch exhibit reduced proliferation and migration compared to those in static culture ^25,26^. We also observed reduced proliferation in 3D compared to 2D culture (Figure S8), emphasizing intrinsic differences in mechanosensing in 2D and 3D model systems.

Our findings of stretch-inhibition of proliferation and invasion further align with a recent study which provides compelling evidence that physiological mechanical load suppresses cancer cell proliferation both *in vitro*, using cyclically contracting cardiac tissue models (cardiomyocyte-laden 3D fibrin gels), and *in vivo* in the beating hearts of mice ^96^. In contrast, mechanical unloading promoted cancer cell proliferation and tumor size *in vivo*, suggesting that the physiological mechanical environment of the heart inherently limits tumor growth. Together with our observation that static conditions promote greater cell proliferation and invasion relative to cyclic stretch, these studies support an emerging framework in which dynamic mechanical loading broadly inhibits invasive and proliferative cell behaviors. This convergence extends the relevance of our findings beyond tissue engineering and into mechanobiological mechanisms of cancer progression and metastasis.

The stretch-induced reduction in cell proliferation we observed may reflect, in part, an elevation of cell-generated tension. Cell tension regulates proliferation through FAK-dependent ^26,97^ and ROCK-dependent signaling pathways ^98^, and this effect is consistent with our finding that blebbistatin, which reduces cell tension, partially rescues the stretch-induced decrease in invasion (Figure 6), thereby supporting a mechanistic relationship between tension and proliferation. However, the combined effects of tension and proliferation account for only moderate inhibition and therefore do not fully explain the profound reduction in invasion observed with stretch. This disparity suggests the involvement of additional stretch-responsive mechanisms such as changes in matrix remodeling ^99,100^ and apoptosis ^32,101^ which warrant further investigation.

### Implications of stretch-inhibition on TEHV repopulation in vivo

In the context of TEHVs, our finding that stretch reduces cell invasion suggests that host-mediated cell repopulation of implanted TEHVs may be less efficient under dynamically loaded conditions. Supporting this, recent work has proposed that the slow recellularization observed in TEHVs may result from the high-amplitude cyclic bending experienced by the valve leaflets ^102^. Although the prevailing paradigm favors implanting cell-free scaffolds that rely on host-driven repopulation *in situ* ^3,5^, our results raise the possibility that static pre-recellularization *in vitro* may enhance scaffold repopulation. We observed substantial cell invasion (>100 μm) within just two days under static conditions indicating that significant cellular infiltration can occur rapidly in the absence of stretch. This finding suggests an alternative strategy in which patient-derived host cells could be seeded on TEHVs and cultured under static culture conditions during the preoperative period prior to implantation. Notably, this strategy has shown clinical success with allografts; e.g., peripheral mononuclear cells isolated from a patient’s blood were reseeded into a decellularized pulmonary valve allograft and cultured *in vitro* for a month prior to implantation in children, resulting in successful reintegration after 3.5 years ^103^. However, TEHVs are typically more porous than allografts ^104^, which suggests that sufficient recellularization could be achieved over a substantially shorter culture period. Pre-seeding TEHVs *in vitro* may therefore accelerate initial cell repopulation and improve tissue integration after implantation, ultimately supporting the development of more integrated, living valve replacements that can grow with the patient.

### Conclusions

In this work, we demonstrate that cyclic stretch profoundly inhibits invasion of cells into surrounding 3D ECM. The direction of uniaxial stretch does not influence the direction of invasion suggesting the reduction is due to an active cell response rather than a passive contact guidance-or stretch-induced matrix stiffening-based mechanism. Cell tension appears to play a role in the reduction of invasion as cell types that are highly contractile experience less disruption of invasion than cell types that are less contractile, and inhibition of myosin IIA partially rescues invasion from stretched spheroids. Reduced cell proliferation also contributes to the stretch-induced inhibition of invasion, potentially through tension-dependent, actomyosin-mediated signaling pathways. Additional mechanisms remain to be explored in future studies, including stretch-induced modulation of matrix remodeling and apoptosis. Because many tissues are inherently dynamic (e.g., heart valves, myocardium, and blood vessels) and because cell invasion and migration are central to both tissue homeostasis and disease, a deeper understanding of how stretch regulates cell invasion in 3D systems is essential. In this context, our work provides insight into how cyclic stretch influences scaffold repopulation in decellularized tissue engineered heart valves while also offering broader relevance to other mechanically driven physiological and pathological processes, such as wound healing and cancer.

## NON-STANDARD ABBREVIATIONS AND ACRONYMS

TEHV: tissue-engineered heart valves
ECM: extracellular matrix
SMC: smooth muscle cells
VIC: valvular interstitial cells
HDF: human dermal fibroblast cells
DOF: depth of field
LPA: Lipoprotein(a)
TGFβ: Transforming growth factor beta

## ACKNOWLEDGEMENTS

We thank Ellie Ungashick, Madison Silva, Delaney Mohl, Temya Jackson Long, Grace Jolin and Kevin Piskorowski for their assistance conducting experiments with collagen and fibrin hydrogels. We also thank Dr. Qi Wen and Pengbo Wang for their assistance with conducting TFM experiments and TFM image analysis. The graphical abstract was made with the assistance of Illustrae software.

## NOVELTY AND SIGNIFICANCE

### What is known?

- Effective *in vivo* cell repopulation of tissue engineered heart valves remains a challenge for the aortic valve, likely driven in part by the dynamic mechanical environment of the valve.
- In 2D systems, the effects of cyclic stretch on cell migration and related behaviors are highly context-dependent, increasing or decreasing depending on stretch magnitude, duration and frequency.
- In 3D scaffolds, stiff mechanical boundary conditions promote directional cell invasion along a defined mechanical axis, yet the effects of dynamic stretch on invasion have not been systematically studied in 3D.

### What new information does this article contribute?

- Cyclic stretch consistently suppresses cell invasion in 3D environments.
- Cyclic stretch reduces cell proliferation, and pharmacologic inhibition of myosin IIA partially rescues invasion with stretch.
- The inhibitory effect of stretch on invasion is independent of stretch direction.

This study provides experimental evidence that cyclic stretch suppresses cell invasion in 3D scaffolds, revealing a previously underexplored inhibitory role for dynamic mechanical loading. We demonstrate that this effect is mediated, in part, by actomyosin-dependent mechanotransduction, as cyclic stretch reduces cell proliferation, and inhibition of myosin IIA partially rescues invasion with stretch. These findings suggest that the dynamic mechanical environment of the heart valve may intrinsically limit host cell infiltration and repopulation of tissue engineered heart valves. Accordingly, strategies incorporating initial static culture prior to implantation may enhance recellularization outcomes. More broadly, this work identifies dynamic mechanical loading as a regulator of cell invasion in 3D environments, with implications for cardiovascular tissue engineering and mechanobiological processes relevant to disease.

